## Supplementary data, tables and figures for "Fungal antigenic variation using mosaicism and reassortment of subtelomeric genes’ repertoires, potentially mediated by DNA triplexes"

### Supplementary data, figures, and tables, Meier et al.

### Supplementary data

##### 1. Reproducibility of the determination of the repertoires

In order to investigate the reproducibility of the whole methodology, eight samples were analysed twice in independent experiments using identical conditions and protocol, except for the DNA extraction that was performed only once. The alleles identified and their abundance were highly similar in the duplicates of four complete repertoires with 73 to 98% of alleles in common (Table S5, Fig. S3a). Moreover, all alleles identified in only one of the duplicates were low abundant (less than 1% of all reads composing the repertoire, gray and light green lines in Fig. S3a). These alleles were presumably not amplified in one duplicate. This can be explained by the stochastic variation in the number of copies of the low abundant genes used as template in the PCR amplification. In other words, low abundant alleles are not amplified each time in repeated PCRs.

As far as the expressed repertoires are concerned, there were more differences between the duplicates than for the complete repertoires with only 33 to 53% of alleles in common. For two samples (BE1 and LA1), most alleles (97 to 100%) that were not present in both duplicates had an abundance lower than 1%, as observed for the complete repertoires (Table S5). On the other hand, the values for samples LA3 and LA4 were only 33 and 50%, respectively. This might result from the lower number of alleles in these two latter samples (14 to 22 versus 82 to 140 in BE1 and LA1). The reduced reproducibility for the expressed repertoires might be due to the greater variation in the allele abundances than in the complete repertoires (see Results, section “Abundance of the *msg*-I alleles in the patients”), a phenomenon that could be increased when the number of alleles present decreases.

#### 2. Confirmation of the allele abundance within patients using amplicon subcloning

The last step of the bioinformatics pipeline provided the abundance of each allele in each sample in percentage of all reads composing the repertoire. A wet-lab approach was used to verify these abundance values. Amplicons from the expressed repertoire of patients LA7, BR3 and SE3 used for PacBio CCS were subcloned using the TOPO cloning kit (Invitrogen). These samples were selected because the low numbers of alleles composing their expressed repertoires facilitates the estimation of abundances by subcloning. Despite the low accuracy of the subcloning approach due to the small number of subclones analysed (8 to 19), the abundances obtained are consistent with those observed using Pacbio CCS followed by the bioinformatics pipeline (Table S6).

##### 3. Search of duplicated fragments within the subtelomeres of a single strain

We searched for all duplicated fragments of size ≥ 100 bps present within the 10 representative subtelomeres assembled from a single strain (Fig. S6). We used BLASTn comparisons of the 35 genes, 15 pseudogenes, and 50 intergenic spaces covering integrally these subtelomeres to the *P. jirovecii* PacBio genome assembly. The significant hits obtained for each gene corresponded to genes and pseudogenes of the same family, whereas pseudogenes generated few hits. The hits obtained for the intergenic spaces were intergenic spaces of the subtelomeres, which were upstream of genes or pseudogenes of the same family as for the query. However, families II and III constituted an exception because some of their upstream sequences produced reciprocal hits with a low score or coverage of the query, which was consistent with the observations made in the section “Sequences flanking the *msg* genes” of the Results. Visual inspection of all alignments of the queries with their hits identified 61 and 38 duplicated fragments involving respectively 14 genes, 5 pseudogenes and 17 intergenic spaces, with a length up to 1142 bps (Table S8). We did not detect any duplicated fragments between the genes and intergenic spaces of family VI, nor between those of families II and III. Alignments of a number of mosaic genes, for example *msg*-I no. 45 and 94 (Fig. S10), suggested that the regions between the shared fragments presented a sequence identity close to those observed on average between the members of each *msg* family (66 to 83%) [1]. The presence of at least one of these duplicated fragments suggests that 22 to 100% of genes, pseudogenes, or intergenic spaces of families I to V are mosaic. These proportions are in fair agreement with those we previously observed [1] for genes and pseudogenes, including the absence of mosaicism in family VI (Table S8). Five out of the 15 pseudogenes investigated were also concerned by the phenomenon. These observations confirm the mosaicism of *msg* genes and pseudogenes, and reveal that the intergenic spaces are also concerned.

The 99 duplicated fragments detected are represented on the 10 representative subtelomeres that we analysed (Fig. S6), as well on the 27 supplementary subtelomeres (Fig. S7). Inspection of Fig. S6 revealed the following features:

(i) Most subtelomeres share fragments with many other subtelomeres.

(ii) Some genes and their upstream space presented many duplicated fragments (for example, gene *msg*-II no.13 in subtelomere 26, and *msg*-III no. 53 in subtelomere 74). Sequence alignments revealed that these fragments sometimes overlap or are identical. This is also revealed by the locations of the shared fragments within the query (Tables S7c and S7d). The important variation of the latter locations suggest that no hotspots of recombination are not present along the *msg* genes.

(iii) The entirety of the partial *msg*-I gene no. 52 at the end of subtelomere 74 (Fig. S6) is duplicated in subtelomere 95 (subtelomere/contig 95, gene no. 61, Fig. S7b), suggesting a whole gene duplication.

**4. Specificity of the CRJE sequence for *P. jirovecii***

Because of the peculiarity of the CRJE, we wondered if it was specific to *P. jirovecii* by using it as query in a BLASTn analysis against the whole nucleotide collection (nr/nt). The full CRJE sequence of 33 bps is present only in *P. jirovecii*, and only at the beginning of *msg*-I genes. Nevertheless, the 25 bps sequence covering the mirror repeats, *i.e.* positions 2 to 26 in Fig. 7a, was present in few copies and few other organisms (2 copies in 1 bacterial species, and 1 to 4 copies in 7 different butterfly species).

**5. Absence of site-specific recombinase targeting the *P. jirovecii* CRJE sequence**

To investigate if of a recombinase that recognizes specifically the CRJE sequence exists in *P. jirovecii*, we performed extensive homology searches using BLASTn involving site-specific recombinases as bait from various organisms (see Methods). These analyses did not detect any recombinases of interest.

### Supplementary tables

**Table S1.** BAL samples from 24 immunocompromised patients analysed in this study.

| City | Country | Patient code [a] | Collection year | Underlying disease |
|---|---|---|---|---|
| Lausanne | Switzerland | LA1 | 2014 | HIV |
| | | LA2 | 2014 | HIV |
| | | LA3 | 2014 | unknown |
| | | LA4 | 2018 | unknown |
| | | LA5 | 2014 | unknown |
| | | LA6 | 2017 | unknown |
| | | LA7 | 2014 | HIV |
| | | LA8 | 2012 | HIV |
| | | LA9 | 2014 | HIV |
| Bern | Switzerland | BE1 | 2014 | HIV |
| | | BE2 | 2013 | kidney transplant |
| | | BE4 | 2015 | cancer |
| | | BE5 | 2014 | cancer |
| Brest | France | BR1 | 2018 | giant cell arteritis |
| | | BR2 | 2018 | cancer |
| | | BR3 | 2019 | HIV |
| | | BR4 | 2020 | psoriasis (methotrexate) |
| | | BR5 | 2019 | cancer |
| Cincinnati | US | CI1 | 2009 | unknown |
| | | CI5 | 1995 | unknown |
| Seville | Spain | SE1 | 2007 | HIV |
| | | SE2 | 2010 | HIV |
| | | SE3 | 2013 | HIV |
| | | SE4 | 2013 | HIV |

[a] LA, Lausanne. BE, Bern. BR, Brest. CI, Cincinnati. SE, Seville.

**Table S2.** R packages used with the R version 4.1.0 (2021-05-18).

| R package | Version | Description | Reference |
|---|---|---|---|
| BiocManager | 1.30.16 | access bioconductor package repository | 2 |
| biostrings | 2.62.0 | manipulation of biological strings | 3 |
| DECIPHER | 2.22.0 | manage biological sequences | 4 |
| dendextend | 1.15.2 | dendogram manipulation | 5 |
| ggplot2 | 3.3.5 | figures and plots | 6 |
| gplots | 3.1.1 | create heatmaps | 7 |
| gridExtra | 2.3 | arrange plots | 8 |
| pBrackets | 1.0.1 | bracket elements in plot | 9 |
| plotrix | 3.8-2 | plot options | 10 |
| reshape2 | 1.4.4 | reshape data | 11 |
| seqinr | 4.2-8 | manipulation of sequences | 12 |
| stringr | 1.4.0 | string operations | 13 |

**Table S3.** Primers used for the control PCRs specific to given alleles.

| Allele | Name allele PacBio | Present in patients [a] | Primer | Position (nt) | Sequence (5' - 3') |
|---|---|---|---|---|---|
| **A** | CI3c106432837 | LA2, LA8, BE1, CI3 | A-for | 198-224 | CCAAGTTGTAAAGGTAGTAAATGTAGC |
| | | | A-rev | 1909-1929 | TAAACGACTTCGTGTCTTTTG |
| **B** | LA2cd106037614 | LA2, CI3 | B-for | 52-70 | GTCCACCTCTAGTGCAAGC |
| | | | B-rev | 1627-1651 | TTCTTGACATTTCCATTTTTATTAG |
| **C** | LA7c108528291 | LA2, BE1, CI3 | C-for | 57-80 | GAAAAGACATGTGAGAACCTTATG |
| | | | C-rev | 1791-1812 | GCTTTTCTAGTTCTTTTCTTGC |

[a] BE, Bern. CI, Cincinnati. LA, Lausanne.

**Table S4.** Characteristics of the expressed and complete *msg*-I gene repertoires observed in the 24 patients.

| Patient [a] | Number of alleles in expressed repertoire | Number of alleles in complete repertoire | Number of alleles in common between the two repertoires | % expressed in complete (alleles in common / total number in expressed) | % complete in expressed (alleles in common / total number in complete) | Number of *P. jirovecii* strains |
|---|---|---|---|---|---|---|
| **LA1** | 82 | 148 | 72 | 88 | 49 | 3 |
| **LA2** | 54 | 49 | 44 | 81 | 90 | 1 |
| **LA3** | 22 | 79 | 20 | 91 | 25 | 1 |
| **LA4** | 18 | 130 | 14 | 78 | 11 | 3 |
| **LA5** | 21 | 144 | 19 | 90 | 13 | 2 |
| **LA6** | 91 | 116 | 82 | 90 | 71 | 5 |
| **LA7** | 5 | 59 | 5 | 100 | 8 | 1 |
| **LA8** | 59 | 105 | 51 | 86 | 49 | 1 |
| **LA9** | 7 | 54 | 7 | 100 | 13 | 1 |
| **BE1** | 90 | 185 | 59 | 66 | 32 | 4 |
| **BE2** | 63 | 56 | 56 | 89 | 100 | 1 |
| **BE4** | 82 | 113 | 79 | 96 | 70 | 3 |
| **BE5** | 18 | 118 | 9 | 50 | 8 | 5 |
| **BR1** | 3 | 83 | 3 | 100 | 4 | 3 |
| **BR2** | 2 | 96 | 2 | 100 | 2 | 3 |
| **BR3** | 5 | 128 | 2 | 40 | 2 | 3 |
| **BR4** | 9 | 96 | 5 | 56 | 5 | 2 |
| **BR5** | 18 | 170 | 17 | 94 | 10 | 4 |
| **CI1** | 108 | 140 | 100 | 93 | 71 | 4 |
| **CI5** | 45 | 44 | 32 | 71 | 73 | 1 |
| **SE1** | 41 | 61 | 37 | 90 | 61 | 1 |
| **SE2** | 12 | 115 | 10 | 83 | 9 | 1 |
| **SE3** | 8 | 77 | 8 | 100 | 10 | 3 |
| **SE4** | 17 | 139 | 17 | 100 | 12 | 3 |

[a] LA, Lausanne. BE, Bern. BR, Brest. CI, Cincinnati. SE, Seville.

**Table S5.** Duplicated analyses of eight samples.

| Repertoire | Patient [a] | Number alleles | | | Common alleles in duplicates | | % of different alleles with abundance <1% |
|---|---|---|---|---|---|---|---|
| | | duplicate 1 | duplicate 2 | Total distinct | number | % (alleles in common / total number distinct alleles) | |
| complete | LA2 | 49 | 48 | 49 | 48 | 98 | 100 |
| | LA1 | 149 | 134 | 154 | 129 | 84 | 100 |
| | BE1 | 185 | 211 | 212 | 184 | 87 | 100 |
| | LA3 | 79 | 80 | 92 | 67 | 73 | 100 |
| expressed | BE1 | 90 | 121 | 138 | 73 | 53 | 100 |
| | LA1 | 82 | 140 | 148 | 74 | 50 | 97 |
| | LA3 | 22 | 19 | 28 | 13 | 46 | 33 |
| | LA4 | 18 | 14 | 24 | 8 | 33 | 50 |

[a] LA, Lausanne. BE, Bern.

**Table S6.** Abundance of alleles determined using PacBio CCS and subcloning.

| Patient [a] | Allele name [b] | Abundance PacBio (%) | Subcloning | |
|---|---|---|---|---|
| | | | Abundance (%) | Nb of clones |
| BR3 | BR3u100403135 | 33 | 74 | 14 |
| | BR3u100532918 | 32 | 0 | 0 |
| | BR3u103089526 | 24 | 21 | 4 |
| | BR3u100992345 | 10 | 5 | 1 |
| | BR3u119802438 | 2 | 0 | 0 |
| LA7 | LA7u47056848 | 72 | 60 | 6 |
| | LA7u100272069 | 9 | 0 | 0 |
| | LA7u10029856 | 8 | 10 | 1 |
| | LA7u101910862 | 7 | 20 | 2 |
| | LA7u100074959 | 4 | 10 | 1 |
| SE3 | SE3u100009239 | 35 | 38 | 3 |
| | SE3u100271002 | 14 | 13 | 1 |
| | BR1u102369376 [b] | 13 | 13 | 1 |
| | SE3u100008413 | 11 | 13 | 1 |
| | CI1u100600372 [b] | 9 | 0 | 0 |
| | SE3u117048602 | 8 | 25 | 2 |
| | SE3u105186188 | 7 | 0 | 0 |
| | SE3u110953735 | 4 | 0 | 0 |

[a] LA, Lausanne. BR, Brest. CI, Cincinnati. SE, Seville.

[b] The name of these alleles results from their high abundance in samples BR1 or CI1.

**Table S7.** See separate excel file.

**Table S8.** Duplicated fragments ≥ 100 bps within the10 *P. jirovecii* representative subtelomeres from a single strain [a].

| | *msg* family | No. of genes or intergenic space upstream of genes (of which pseudogenes): used as query | No. of genes or intergenic space upstream of genes (of which pseudogenes): with duplicated fragments ≥ 100 bps | Total no. of duplicated fragments ≥ 100 bps detected [b] (of which between genes and pseudogenes) | Mean size (bps) of duplicated fragments (range) | % potential mosaic genes or intergenic space (pseudogenes included) | % potential mosaic genes in reference 1[c] |
|---|---|---|---|---|---|---|---|
| Genes and pseudogenes | I | 20 (8) | 5 (2) | 7 (2) | 513 (109-670+1142) [d] | 25 | 42 |
| | II | 9 (3) | 5 (0) | 32 (0) | 249 (117-710) | 56 | 28 |
| | III | 5 (0) | 3 (0) | 12 (0) | 120 (100-153) | 60 | 40 |
| | IV | 2 (2) | 2 (2) | 4 (4) | 165 (113-221) | 100 | 22 |
| | V | 7 (1) | 4 (1) | 7 (1) | 183 (102-363) | 57 | 7 |
| | VI | 4 (0) | 0 (0) | 0 (0) | 0 | 0 | 0 |
| | outlier | 3 (1) | 0 (0) | 0 (0) | 0 | 0 | - |
| | **Total** | **50** (15) | **19** (5) | **61** (7) | | | |
| Intergenic spaces | I | 20 (8) | 9 (1) | 12 (1) | 148 (105-224) | 45 | - |
| | II | 9 (3) | 2 (0) | 6 (0) | 314 (131-787) | 22 | - |
| | III | 5 (0) | 2 (0) | 6 (0) | 233 (132-320) | 40 | - |
| | IV | 2 (2) | 1 (1) | 2 (2) | 129 (115, 142) | 50 | - |
| | V | 7 (1) | 3 (1) | 12 (3) | 178 (115-334) | 43 | - |
| | VI | 4 (0) | 0 (0) | 0 (0) | 0 | 0 | - |
| | outlier | 3 (1) | 0 (0) | 0 (0) | 0 | 0 | - |
| | **Total** | **50** (15) | **17** (3) | **38** (6) | | | |

**Footnotes of Table S8 :**

[a] All *msg* genes, pseudogenes, and intergenic spaces composing the 10 representative subtelomeres shown in Fig. S6 were used as query in BLASTn analyses (search of somewhat similar sequences) against the PacBio *P. jirovecii* genome assembly [1]. Each upstream intergenic space extended up to the end of the gene located upstream (all genes are oriented towards the telomere within the subtelomeres, see Fig. S6). Visual inspection of all alignments of the query with the significant hits identified the duplicated fragments. Outlier genes are those that could not be attributed to one of the six *msg* families [1].

[b] The duplicated fragments detected twice because of reciprocal BLASTn analyses were counted only once.

[c] Using various numerical bioinformatics tools to detect recombination events between *msg* genes and pseudogenes [1].

[d] The duplicated fragment of 1142 bps corresponds to the entirety of the partial gene *msg*-I no. 52 at the end of contig 74 (Fig. S6).

### Supplementary figures

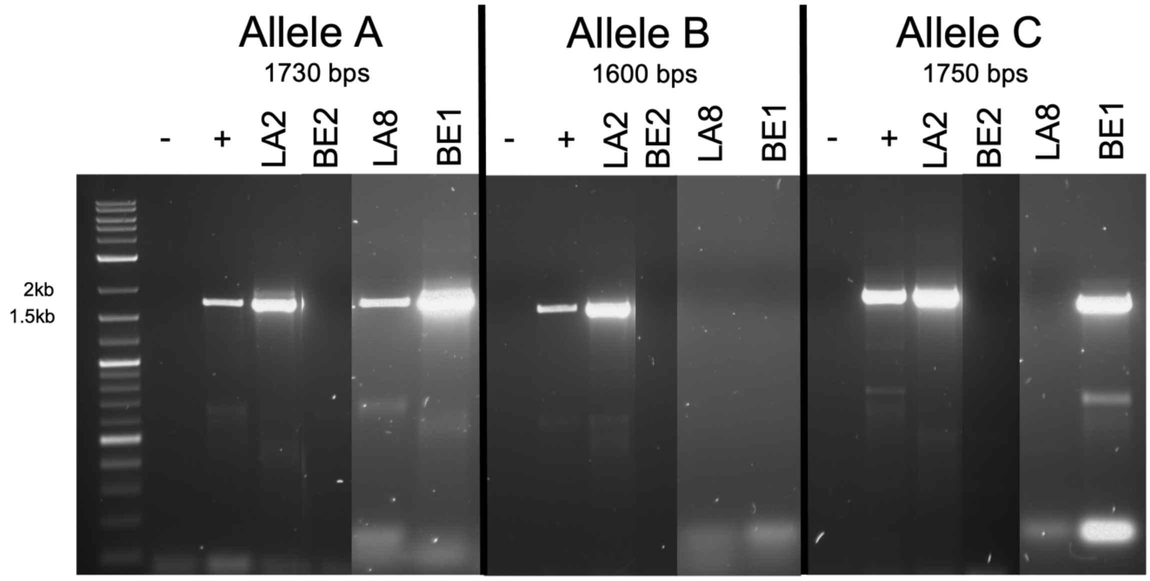

**Figure S1.** Analysis in agarose gel of the PCRs using primers specific to fragments of *msg*-I alleles A, B and C. The size of the PCR product from each allele is indicated. The positive control (+) is randomly amplified DNA of patient LA2 that contains all three alleles. The primers used and alleles are described in Table S3.

10xT

GAAAATTCAGCTTAAACACTTCCCTAGTGTTTTAGCATTTTTCAAACATCTGTGAATTTTTT

6xT

TTTTGTTTGGCGAGGAGCTGGCTTTTTTGCTTGCCTCGCCAAAGGTGTTTATTTTTAAAATT

TTAAATTGAATTTCAGTTTTAGAATTTTTTAAAAACTTTCAACAATGGATCTCTTGGCTCTC

GCGTCGATGAAGAACGTGGCAAAATGCGATAAGTAGTGTGAATTGCAGAATTTAGTGAATCA

TCGAATTTTTGAACGCATCTTGCGCTCCTTAGTATTCTAGGGAGCATGCCTGTTTGAGCGTT

5xT 5xA 3xA

ATTTTTAAGTTCCTTTTTTCAAGCAGAAAAAAGGGGATTGGGCTTTGCAAATATAATTAGAA

TAAAATAATTATATGCATGCTAGTCTGAAATTCAAAAGTAGCTTTTTTTCTTTGCCTAGTGT

4xT

CGTAAAAATTCGCTGGGAAAGAAGGAAAAAAGCTTTTATAAATACAAGAATT

**Figure S2.** *P. jirovecii* ITS1-5.8S-ITS2 sequence (JQ365709.1). Highlighted in yellow are the six homopolymers that were homogenized in the present study, *i.e.* they were replaced in all sequences by the number of nucleotides indicated in red.

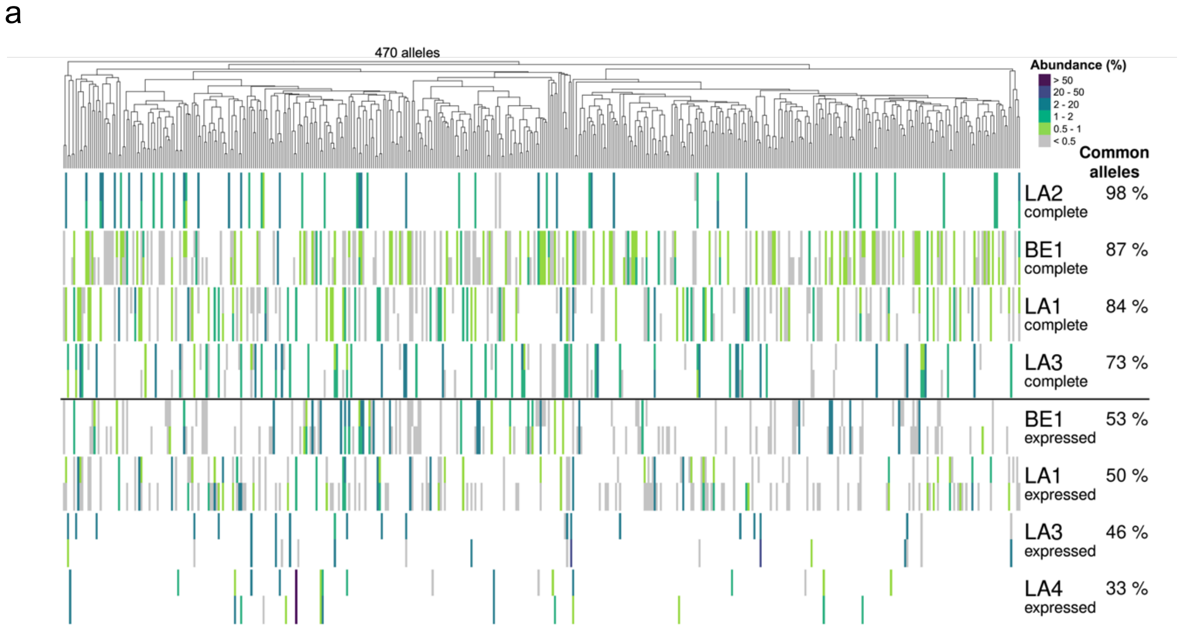

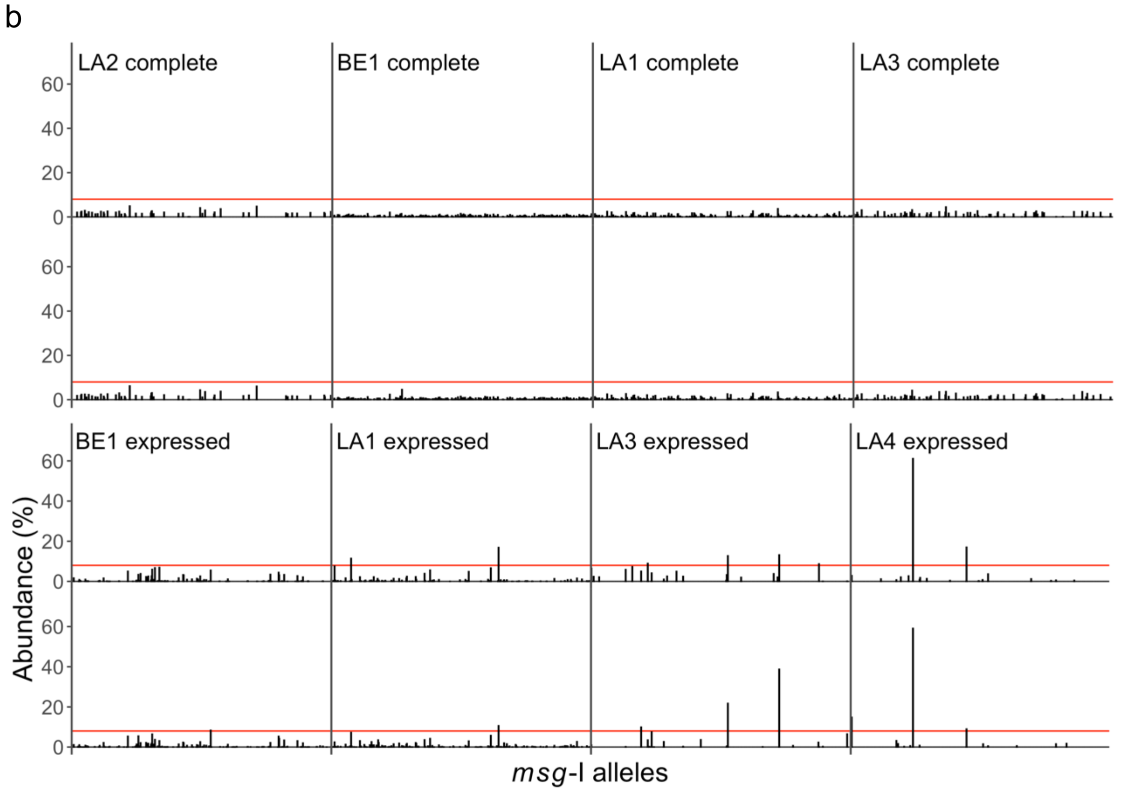

**Figure S3.** Duplicate analyses to evaluate the reproducibility of the whole methodology. LA, Lausanne. BE, Bern.

a. Composition of the complete and expressed *msg*-I repertoires observed. Each vertical line of the heatmap represents an allele present in the given repertoire with the color figuring its abundance in % of all reads composing the repertoire, as indicated at the top right of the figure. The 470 alleles observed were sorted using hierarchical classification trees of the multiple alignments of the allele sequences (Fitch distance, average linkage).The percentage of common alleles between the duplicates are indicated next to the patient's name.

b. Abundance of the alleles within the complete and expressed repertoires (see Results, section "Abundance of the *msg*-I alleles in the patients"). The alleles are sorted using the same tree as in panel a. The red lines indicate an abundance of 8.0%.

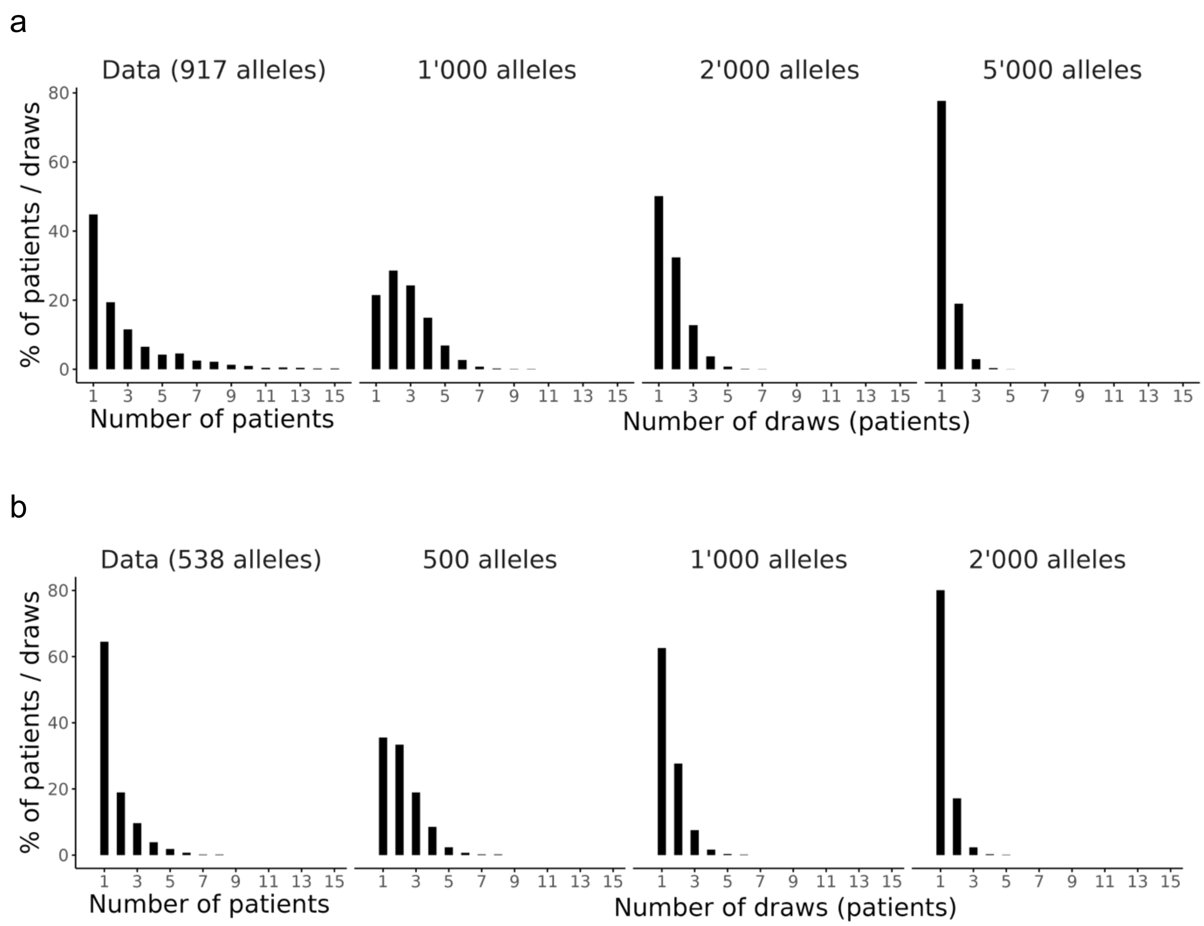

**Figure S4.** Comparison of the distribution of the alleles observed in the complete (a) and expressed (b) repertoires of the 24 patients with those obtained by simulating reservoirs of alleles of increasing size (see text). The software R was used.

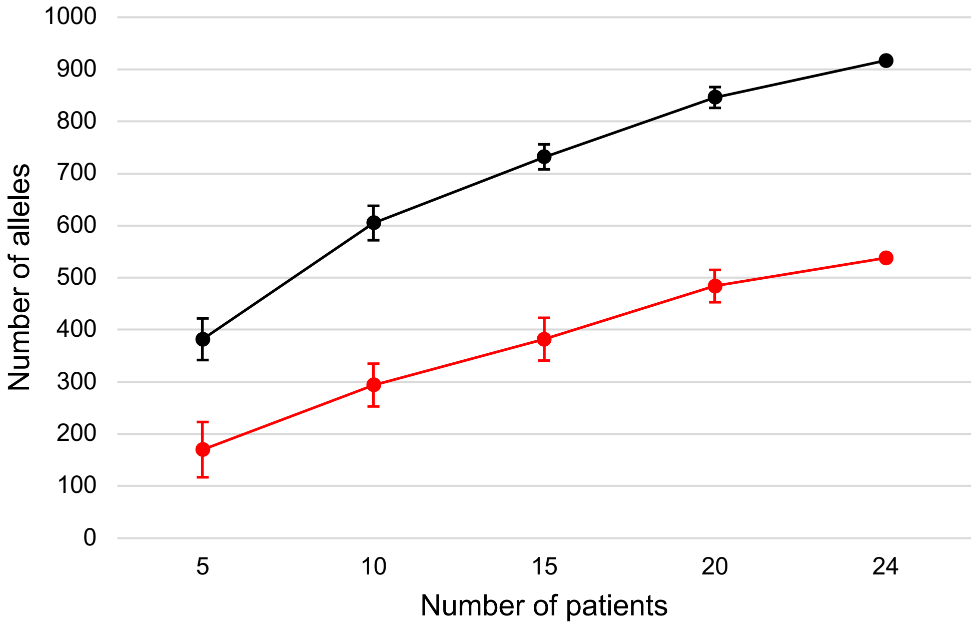

**Figure S5.** Number of alleles observed in the complete (black) and expressed (red) repertoires in function of the simulated number of patients analysed (see text). SD are shown. The software R was used.

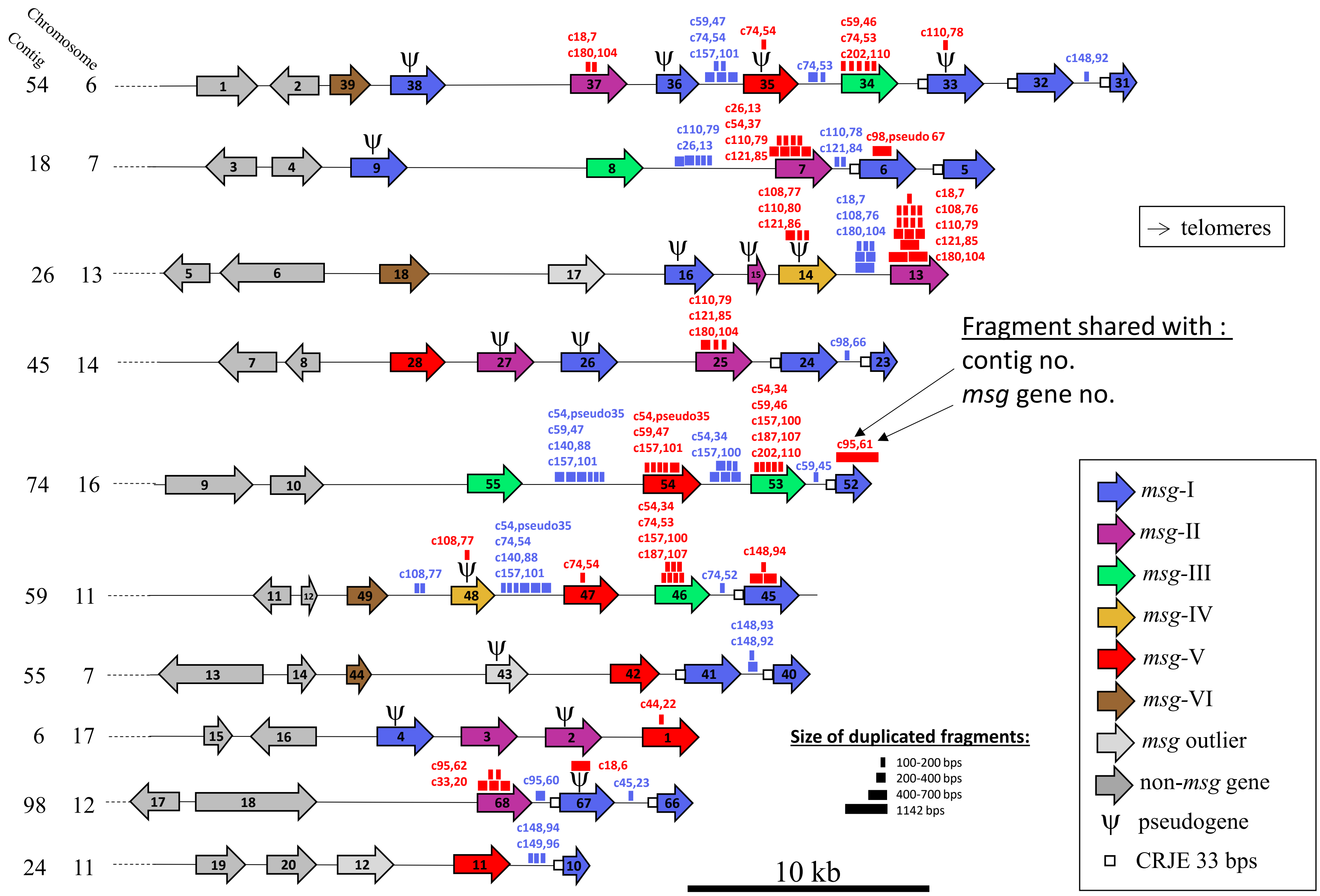

Chromosome
Contig
→ telomeres
Fragment shared with :
contig no.
msg gene no.
Size of duplicated fragments:
100-200 bps
200-400 bps
400-700 bps
1142 bps
10 kb
msg-I
msg-II
msg-III
msg-IV
msg-V
msg-VI
msg outlier
non-msg gene
pseudogene
CRJE 33 bps

**Figure S6.**

**Figure S6.** Ten representative subtelomeres (*i.e.* contigs, see Methods) which genes, pseudogenes, and intergenic spaces were used as queries against the whole genome assembled from a single *P. jirovecii* strain using PacBio sequencing [1]. The symbols represent the size of the duplicated fragments identified within genes and pseudogenes (red symbols), or intergenic spaces (blue symbols). The position of the symbols is not precise. The positions of the duplicated fragments within the query and the subtelomeres are given in Tables S7c and S7d. Adapted from Fig. 3 of reference [1].

a

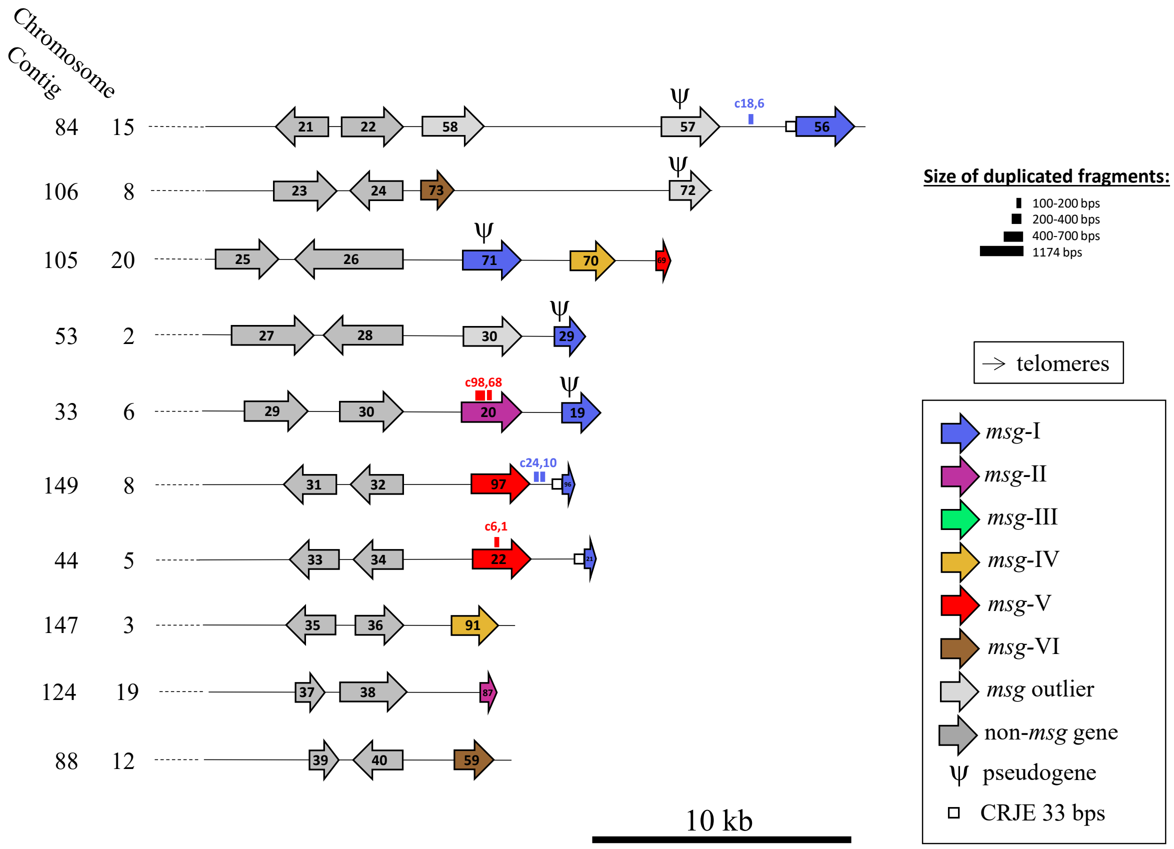
Chromosome
Contig
84 15
106 8
105 20
53 2
33 6
149 8
44 5
147 3
124 19
88 12
21 22 58 57 56
c18,6
23 24 73 72
25 26 71 70
27 28 30 29
29 30 20 19
c98,68
31 32 97
c24,10
33 34 22
c6,1
35 36 91
37 38 87
39 40 59
Ψ
10 kb
Size of duplicated fragments:
100-200 bps
200-400 bps
400-700 bps
1174 bps
→ telomeres
msg-I
msg-II
msg-III
msg-IV
msg-V
msg-VI
msg outlier
non-msg gene
Ψ pseudogene
CRJE 33 bps

**Figure S7.**

b

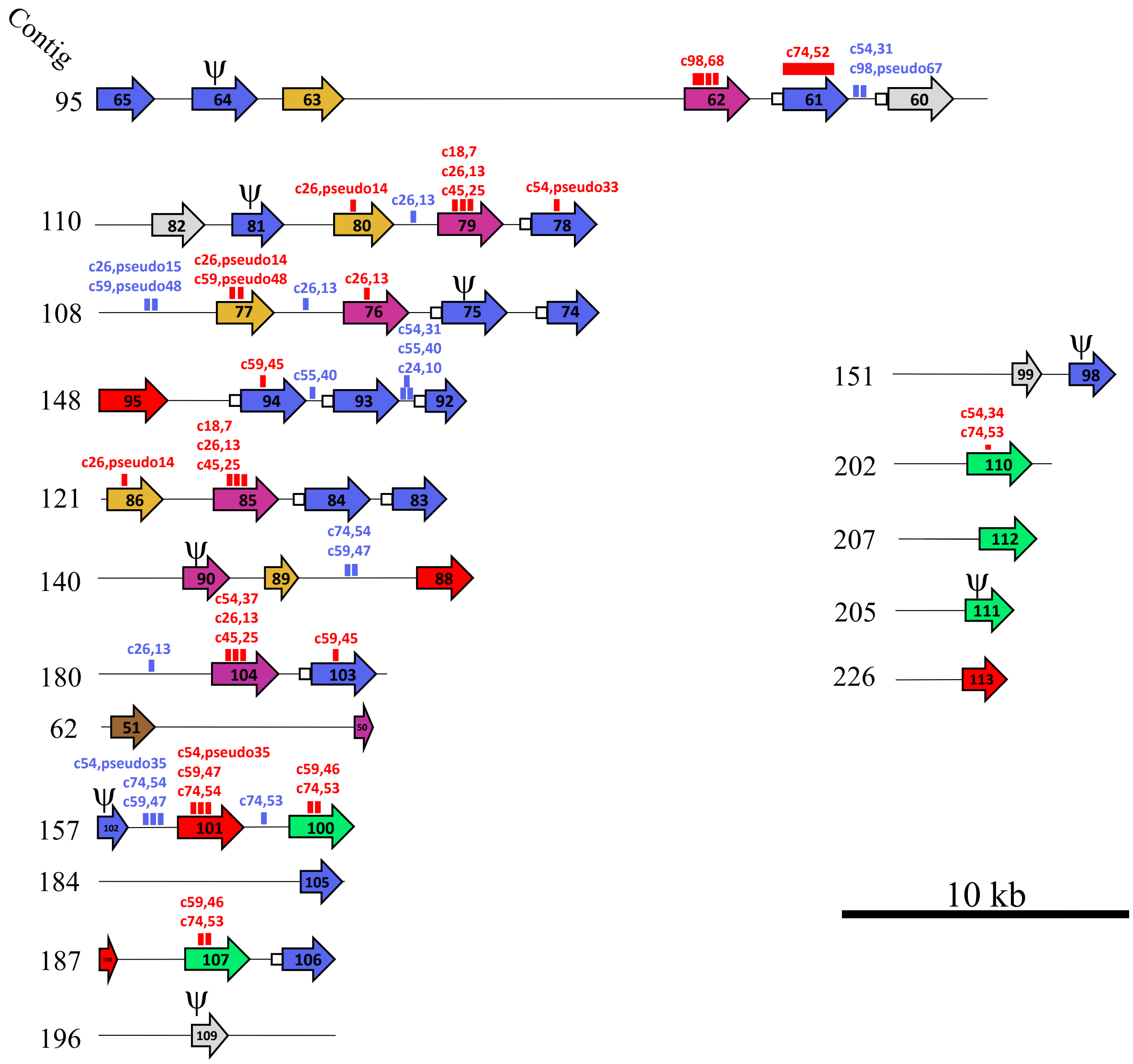

**Figure S7.** Supplementary subtelomeres (*i.e.* contigs, see methods) from the same single strain as the 10 representative ones shown Fig. S6. The symbols represent the size of the duplicated fragments identified within genes and pseudogenes (red symbols), or intergenic spaces (blue symbols). Their positions within the query and the subtelomeres are given in Tables S7c and S7d. Adapted from Fig. S6 of reference [1].

a. 10 subtelomeres harboring genomic genes (in gray) that allowed their attribution to a specific chromosome from reference [14].

b. 17 subtelomeres not harboring genomic genes, preventing their attribution to a chromosome.

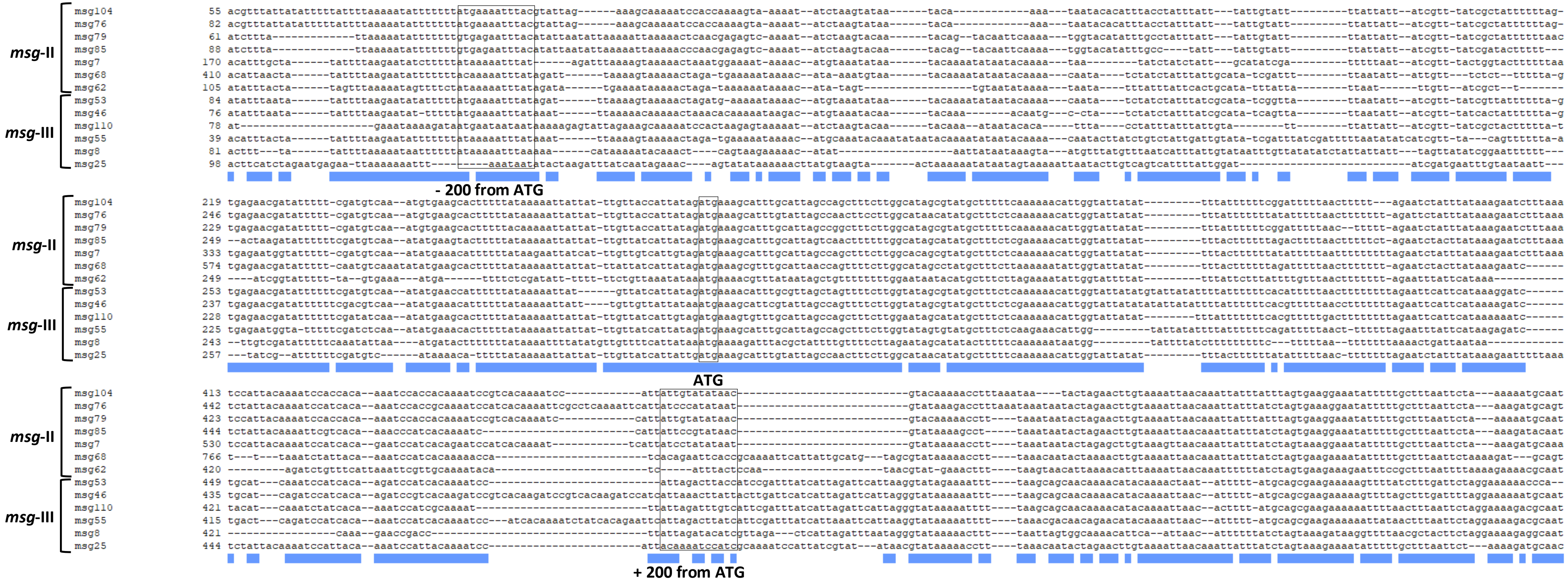

**Figure S8.** Alignments of the region surrounding the ATG of seven *msg*-II and six *msg*-III genes presenting significant similarity in the 200 bps upstream of their CDS. The blue bars underneath indicate areas of significant similarity. Such areas are particularly long around the start codon ATG of the CDS. The ATG start codons and the regions encompassing the – and + 200 bps positions relatively to the ATG of sequences are boxed.

.

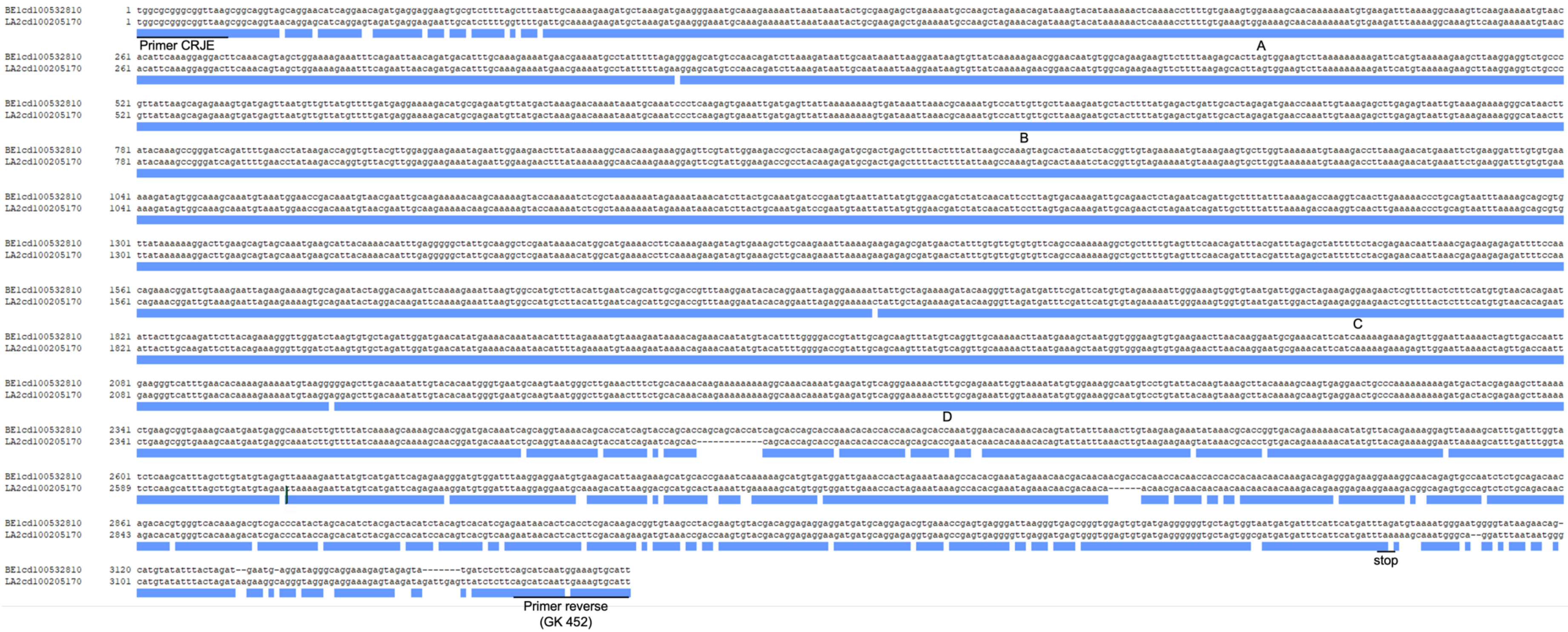

**Figure S9.** Alignment of two mosaic *msg*-I genes identified in this study (the names of the alleles are given on the left). The blue bars indicate areas of full identity. The four fragments of ≥ 100 bps shared are indicated (A to D, respectively of 284, 1335, 420, and 294 bps). Fragment C is shown in red within Figure 6. Clone Manager 9 Professional Edition software version 9.51 (Sci Ed Software LLC) and the alignment format “similarity summary bars” were used.

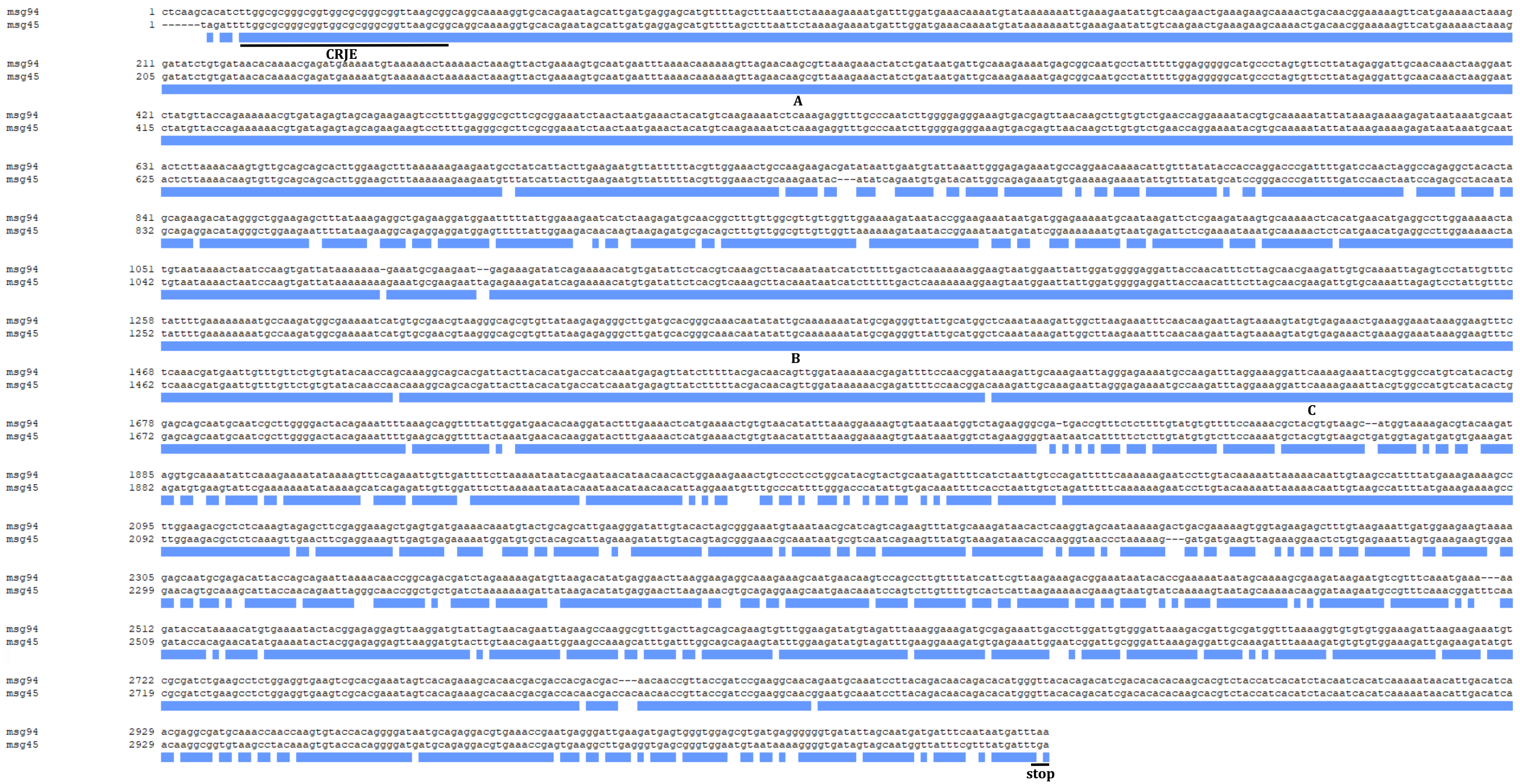

**Figure S10.** Alignment of the mosaic *msg*-I genes no. 45 and 94. The blue bars indicate areas of full identity. The three fragments of ≥ 100 bps shared are numbered (respectively 670, 427, and 118 bps). Clone Manager 9 Professional Edition software version 9.51 (Sci Ed Software LLC) and the alignment format “similarity summary bars” were used.

A.

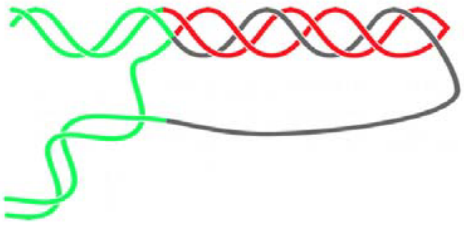

B. **Hoogsteen Triads** **Reverse Hoogsteen Triads**

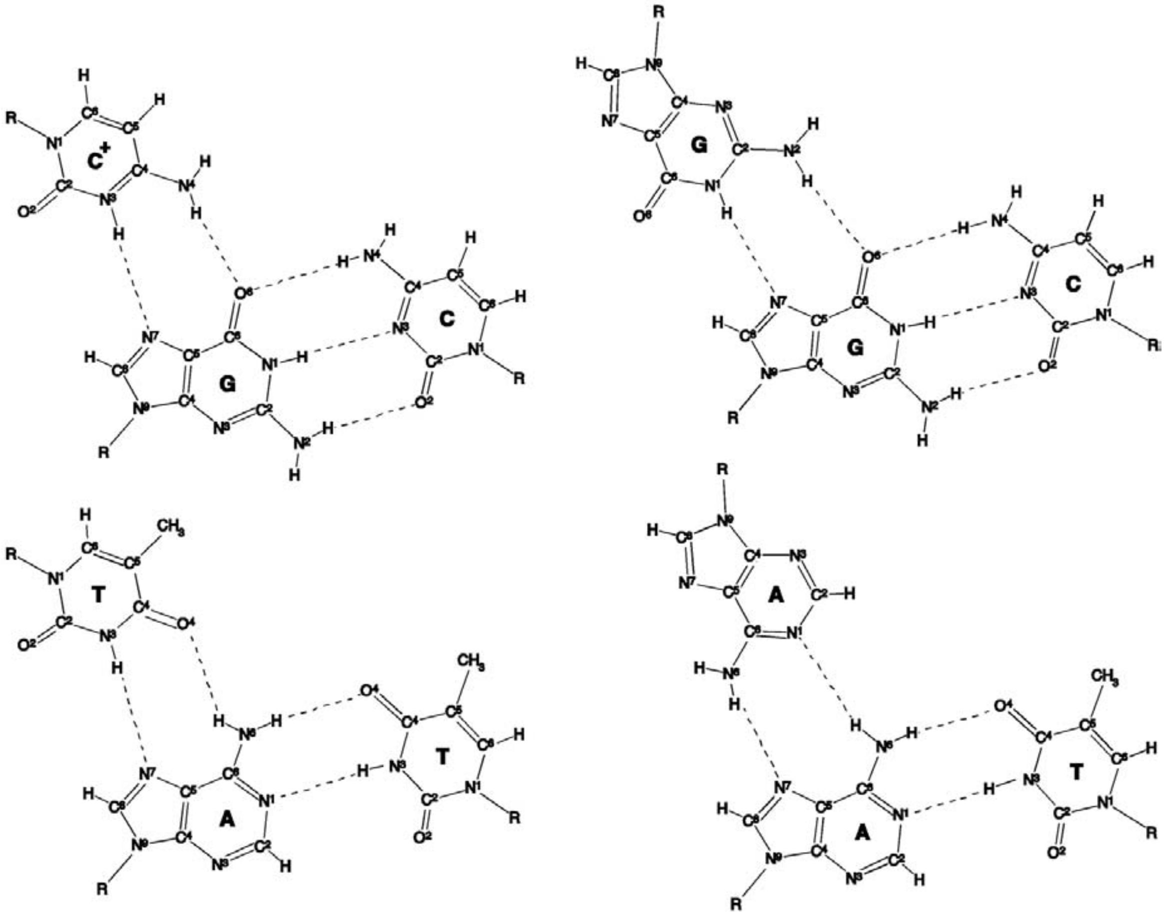

**Figure S11.**

C.

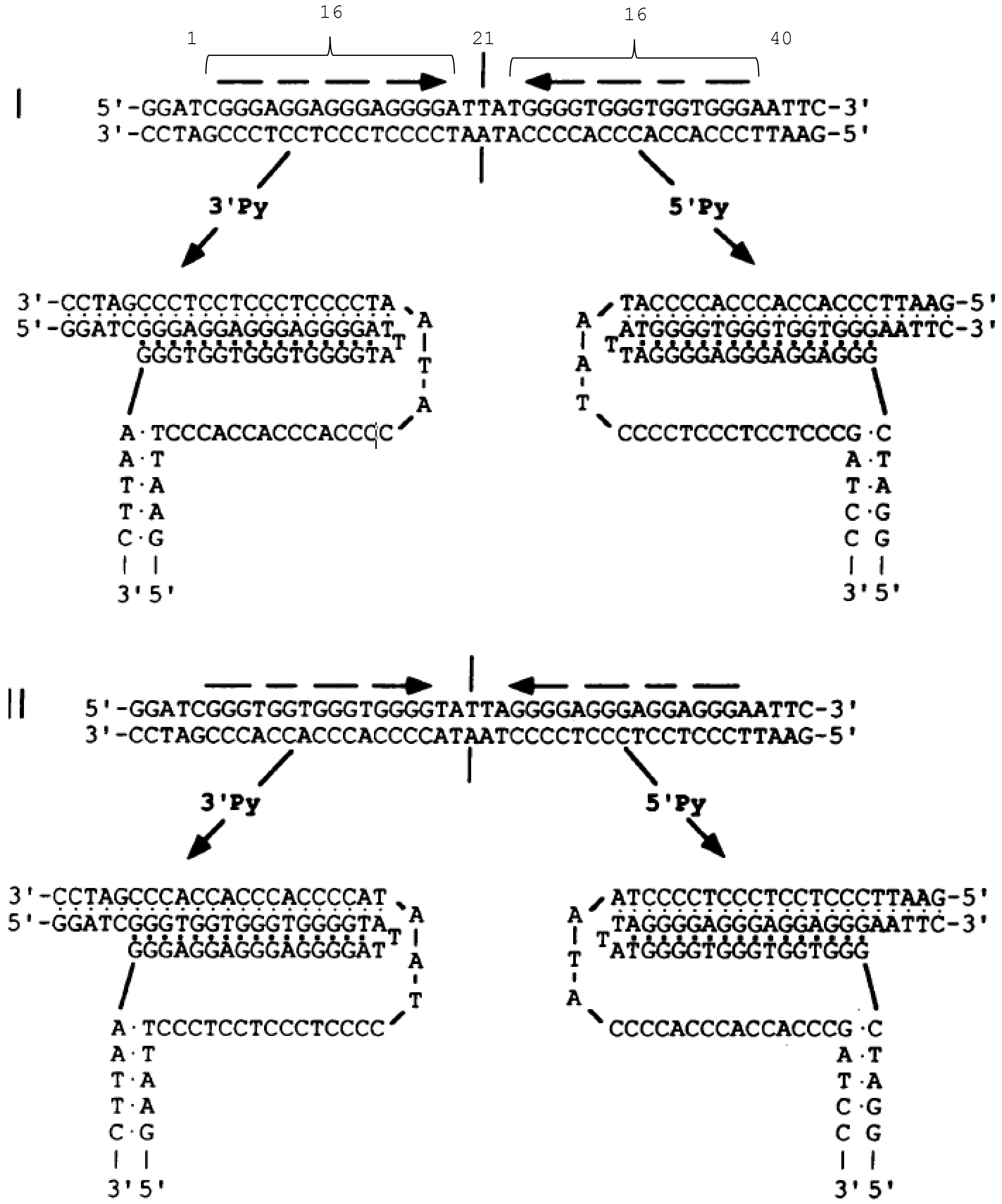
16
16
1
21
40
I
5'-GGATCGGGAGGAGGGAGGGGATTATGGGGTGGGTGGTGGGAATTC-3'
3'-CCTAGCCCTCCTCCCTCCCCTAATACCCCACCCACCACCCTTAAG-5'
3'Py
5'Py
3'-CCTAGCCCTCCTCCCTCCCCTA
5'-GGATCGGGAGGAGGGAGGGGAT
GGGTGGTGGGTGGGGTA
A·TCCCACCACCCACCCC
TACCCCACCCACCACCCTTAAG-5'
ATGGGGTGGGTGGTGGGAATTC-3'
TAGGGGAGGGAGGAGGG
CCCCTCCCTCCTCCCG·C
3'5'
II
5'-GGATCGGGTGGTGGGTGGGGTATTAGGGGAGGGAGGAGGGAATTC-3'
3'-CCTAGCCCACCACCCACCCCATAATCCCCTCCCTCCTCCCTTAAG-5'
3'Py
5'Py
3'-CCTAGCCCACCACCCACCCCAT
5'-GGATCGGGTGGTGGGTGGGGTA
GGGAGGAGGGAGGGGAT
A·TCCCTCCTCCCTCCCC
ATCCCCTCCCTCCTCCCTTAAG-5'
TAGGGGAGGGAGGAGGGAATTC-3'
ATGGGGTGGGTGGTGGG
CCCCACCCACCACCCG·C
3'5'

**Figure S11.**

**Figure S11.** Features of H-DNA and *H-DNA triplexes. A. and B.: Figure 3 and its legend from reference [15]. C.: Figure 1 and its legends from reference [16], with added numbering. All three panels are reproduced with permission from corresponding author and copyright holder S. M. Mirkin.

A. The structure of an intramolecular triplex. The two complementary strands of a homopurine-homopyrimidine repeat are colored in red and gray, while flanking DNA is colored green. The structure is called H-y when the red strand is homopyrimide, and H-r if when it is homopurine. One can see that the red and gray strands in this structure are not linked, *i.e.* formation of H-DNA is topologically equivalent to an unwinding of the entire homopurine-homopyrimidine repeat.

B. H-y form is built from TA*T and CG*$C^+$ triads, in which pyrimidines in the third strand form Hoogsteen hydrogen bonds with the purines of the duplex. H-r form and is built of CG*G and TA*A triads, where purines from the third strand form reverse Hoogsteen hydrogen bonds with the purines in the duplex.

C. Intramolecular triplexes consisting of GG*C and TA*T triads. In both sequences I and II, GC base pairs are arranged as mirror images (shown by arrows; the pseudosymmetry axis is shown by a vertical line), whereas AT base pairs are arranged as inverted repeats. Points, Watson-Crick hydrogen bonds; squares, Hoogsteen hydrogen bonds.
